# Genome-wide prediction of bacterial effectors across six secretion system types using a feature-based supervised learning framework

**DOI:** 10.1101/255604

**Authors:** Andi Dhroso, Samantha Eidson, Dmitry Korkin

## Abstract

Gram-negative bacteria are responsible for hundreds of millions infections worldwide, including the emerging hospital-acquired infections and neglected tropical diseases in the third-world countries. Finding a fast and cheap way to understand the molecular mechanisms behind the bacterial infections is critical for efficient diagnostics and treatment. An important step towards understanding these mechanisms is discovering bacterial effectors, the proteins secreted into the host through one of the six common secretion system types. Unfortunately, current effector prediction methods are designed to specifically target one of three secretion systems, and no accurate “secretion system-agnostic” method is available.

Here, we present PREFFECTOR, a computational feature-based approach to discover effectors in Gram-negative bacteria without prior knowledge on bacterial secretion system(s) or cryptic secretion signals. Our approach was first evaluated using several assessment protocols on a manually curated, balanced dataset of experimentally determined effectors across all six secretion systems as well as non-effector proteins. The evaluation revealed high accuracy of the top performing classifiers in PREFFECTOR, with the small false positive discovery rate across all six secretion systems. Our method was also applied to four bacteria that had limited knowledge on virulence factors or secreted effectors. PREFFECTOR web-server is freely available at: http://korkinlab.org/preffector.

## Introduction

With advancement of genome sequencing and high-throughput methods, it has become clear that the bacterial infection involves a complex interplay of macromolecular interactions between the host proteins and the bacterial effectors, a group of pathogenic proteins [1-3]. Effectors are delivered into the host organism through specialized molecular machinery called the secretion system. Currently, six distinct types of secretion system have been identified for Gram-negative bacteria [4, 5]. Some molecular mechanisms engaged during the secretion of effectors have been characterized [5]. However, the main principles behind targeted selection of effectors among thousands of other bacterial proteins as well as their delivery into the host by the secretion system of each type are far from been fully understood due to the complexity of this question. For instance, it is not uncommon for a bacterial species to secrete dozens, even hundreds of effectors [6, 7] or to harbor more than one secretion system [8-10].

Several families of effectors have been known to carry a translocation signal that is used to recognize bacterial effectors by its secretion system [11, 12]. In some secretion systems this signal is located in the C-termini of the effectors [13-15], and in other systems the signal is found in the N-termini [16, 17]. However, for many effectors no such signal is found, and it has been suggested that the signal can exist anywhere in the protein sequence, and not necessarily in its terminal regions [11, 18]. Knowledge of effectors of a bacterial pathogen is the first step towards efficient design of high-throughput experiments to uncover and mechanistically understand the host-pathogen interaction networks, with the ultimate goal of developing drugs that target key network nodes and connections [19, 20]. Unfortunately, the abundance of effectors, their sequence, structural, and functional diversity, as well as their cryptic recognition signal makes this task extremely challenging.

Recently, several experimental and computational approaches have been developed to identify effectors in Gram-negative bacteria on a large, often whole-genome, scale (Table 1) [21-28]. However, the experimental approaches were designed to find effectors in the genomes of individual bacterial species. Likewise, the computational approaches focus on a single type of secretion system or even a single family of effectors within a secretion system, mainly covering only type III and type IV secretion systems (T3SS and T4SS, respectively). These computational strategies often build on detecting the presence of an explicit translocation signal motif by leveraging supervised learning approaches, such as artificial neural networks (ANN), naïve Bayes classifier, random forest, hidden Markov model (HMM) and support vector machines (SVMs) [24-26, 28] [29]. Alternatively, some supervised learning methods use more “focused” training sets (*e.g.*, a subset of effectors that carry a specific function, or are from specific organisms) [21, 24]. Applying highly accurate but highly focused effector predicting methods to the bacterial genomes for which no secretion system information is available or which could harbor more than one secretion system, could be challenging due to a potentially large number of false positives.

**Table 1.** The current state-of-the-art methods and their reported performance. Each method is designed to predict effectors of a specific type of secretion system. The employed methods include Random Forest (RF), Artificial Neural Network (ANN), Hidden Markov Model (HMM), and Support Vector Machine (SVM).

| *Name* | *Method* | *Secretion System* | *Signal Based* | *Reported Accuracy* | *References* |
| --- | --- | --- | --- | --- | --- |
| Luo *et al* | RF | I | Yes | 88.6% | [21] |
| Luo *et al* | RF | I | No | 98.1% | [21] |
| SSE-AAC | SVM | III | Yes | 98.3% | [64] |
| BPBAac | SVM | III | Yes | 95.3% | [25] |
| T3_MM | Markov Model | III | Yes | 88.2% | [65] |
| BEAN | HMM/SVM | III | No | 78.0% | [26] |
| EffectiveT3 | Naïve Bayes | III | No | 78.0% (AUC) | [22] |
| Sieve | SVM | III | No | 95.0% (AUC) | [66] |
| Yang X *et al* | RF | III | No | 94.3% | [29] |
| T4SEpre family | SVM | IV | Yes | 79.1%-94.6% | [67] |
| T4EffPred | SVM | IV | No | 95.9% | [27] |

In this work, we present an accurate sequence-based approach that draws on machine learning and statistical methods to predict bacterial effectors of Gram-negative bacteria, irrespective of the secretion systems they utilize. Our approach is driven by the following challenges: (1) one needs a “universal” high-throughput method that is capable of detecting effectors of each secretion system, while (2) in general, one does not know the kind of secretion system(s) that the species has and the location or form of the secretion signal.

Our approach, called **Pr**ediction of **Effector**s (PREFFECTOR), builds on previous studies that identified the presence of signals in the effector sequences, without defining the signal explicitly. Specifically, six classifiers, including three SVM and three Random Forest models, have been trained to evaluate three independent hypotheses about the location of a signal motif, followed by comparison of the top performing classifier with several integration schemas that combine the results of all three classifiers (Fig. 1A). When evaluated, PREFFECTOR showed strong performance across all types of secretion systems, and with no prior information about the known effectors of the same genome available, achieving the same maximum of 89% in leave-one-out (LOO) accuracy (Acc), precision (Pre), and recall (Rec) measures (Fig. 1C).

**Figure 1.**
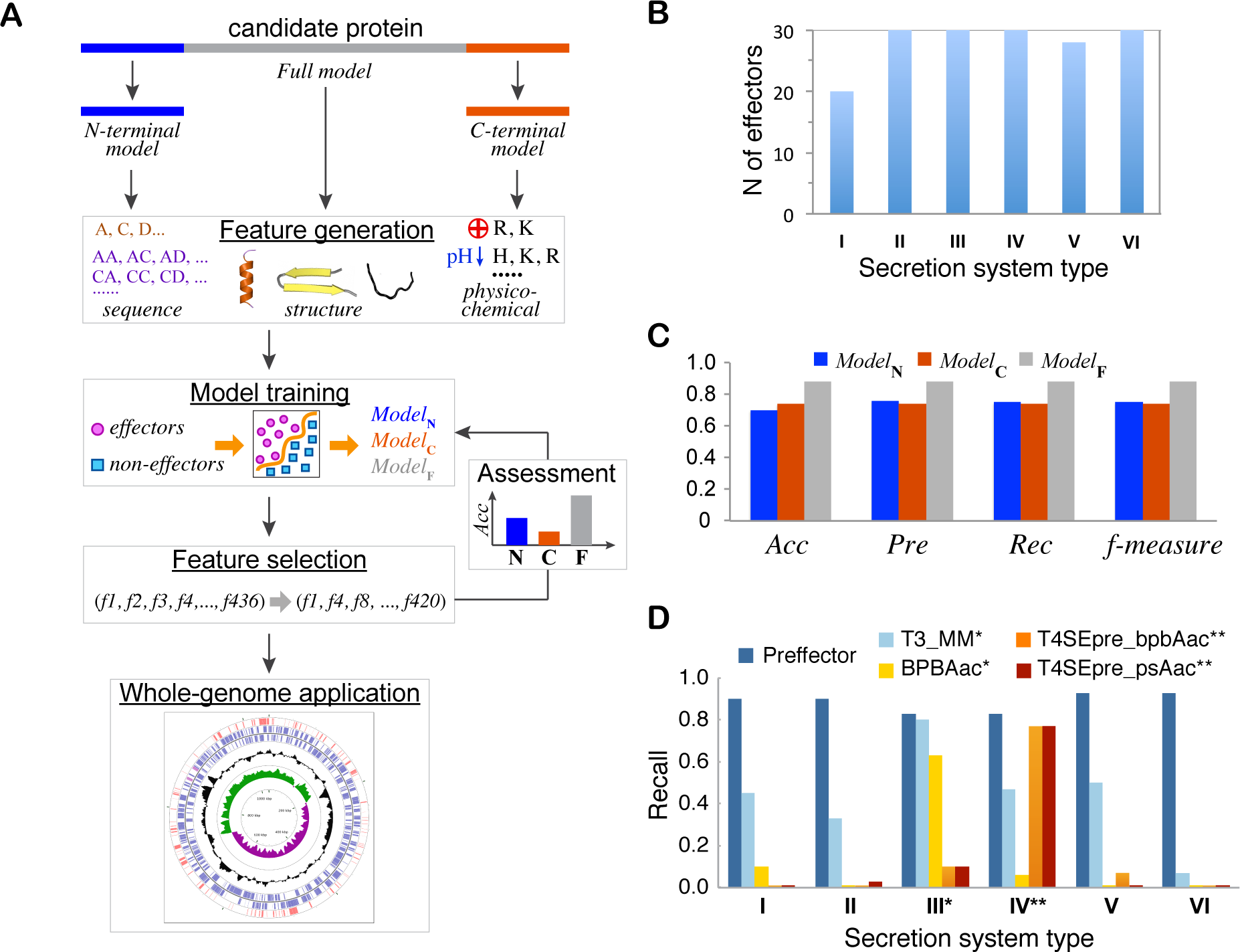
PREFFECTOR and its comparative performance. A. Basic stages of the approach. Three classifier models are designed and independently trained with secretion signals located primarily in the N-terminal region, C-terminal region, or anywhere in the effector protein sequence. Models that rely on the full protein sequence or its N‐ and C-termini are referred to in the figure as *Model*_F_, *Model*_N_, and *Model*_C_, correspondingly. B. The finalized positive set of 168 effectors is manually curated and covers all six secretion systems. C. The most accurate full model (Random Forest classifier) performs evenly across all four assessment measures; it is more accurate than the best performing C-terminal and N-terminal models (both are SVM classifiers with RBF kernels). Abbreviations: Accuracy (Acc), Precision (Pre), and Recall (Rec). D. The assessment of the state-of-the-art methods that are designed to detect effectors of a single secretion system on our data set has showed that none of these methods can be universally used across all six secretion systems. One asterisk (*) corresponds to the methods specialized in predicting T3SS effectors, while two asterisks (**) correspond to the methods specialized in predicting T4SS effectors.

## Methods

Our effector classification approach leverages feature-based supervised learning methods. Each candidate protein is first represented as a 437-dimensional feature vector (Fig. 1A). The calculated features are selected based on the properties found to play an important role in effectors by a number of studies [27, 28]. Next, guided by the current knowledge about the signal locations in the termini of known effectors, three groups of classifiers are designed and independently trained. Each classifier tests a hypothesis that the cryptic secretion signal is located (a) primarily in the N-termini of the effector protein sequence, (b) primarily in its C-termini, or (c) anywhere in the effector sequence. As a result, the feature vectors are calculated using 25 N-terminal residues for the first group of classifiers, 25 C-terminal residues for the second group, and the entire protein sequence for the third group. Each classifier is trained on a balanced non-redundant set of effectors from six secretion system types, collected from the literature and manually curated (Fig. 1B, Supplementary Tables S1, S2).

### A curated dataset of effectors and non-effectors for classification

To train and test the supervised learning classifiers, we first collected a dataset of positive (effectors) and negative (non-effectors) examples. Collecting such a dataset meets three basic challenges. First, to exclude the generalization bias, the positive dataset of effectors should be balanced across all six secretion systems. Unfortunately, while some well-studied secretion system types, such as T3SS, have hundreds of experimentally validated effectors, others, such as type I secretion system (T1SS), have only a couple dozen of confirmed effectors. The secretion system types with the limited number of effectors will therefore constrain the overall size of the balanced positive set. Second, effectors from different species but the same secretion system type often share homology, which is reflected in the high sequence similarity. Thus, it is important to minimize potential bias due to the presence of similar sequences in the dataset. A desirable approach to removing redundancy from the training set is to use a fully automated protocol based on a single threshold. While it can be done for the negative set, for the positive training set one has to select a reasonable trade-off between the maximum coverage for each of the six secretion systems while maintaining the minimal relationship between the effectors within each secretion system. Therefore, while most of the positive training set was obtained using the same automated protocol as the negative training set, we had to further expand the positive set using manual curation (*e.g.,* ensuring that the effectors that share more than 30% sequence similarity had diverse functions and/or came from only remotely related bacterial species). Finally, we note that unlike any of the methods focused on a single secretion system, where a T3SS effector can be correctly labeled as a negative example for a T4SS effector prediction method, our negative set cannot include effectors from any secretion system, requiring careful manual curation of the dataset.

### Supervised feature-based classifier of bacterial effectors

For each protein sequence, we calculate a vector of length-invariant features; the feature vector is then used as an input for the classification models (Table 2). The features are calculated independently for each of the three models. Three categories of features are considered: residue composition, sequence/structure information, and physico-chemical properties of proteins. All features are scaled using basic standardization procedure.

**Table 2.** Three categories of features. The features are calculated independently for each of the three models.

| *Category* | *Feature position in vector* | *Feature Description* | *Dimensions* |
| --- | --- | --- | --- |
| Structure/sequence information | 1 | Length | 1 |
|  | 432-436 | H, C, E, Exposed, Buried | 5 |
| Residue composition | 4-23 | Residue occurrence frequency | 20 |
|  | 32-431 | Dipeptide occurrence frequency | 400 |
| Physico-chemical properties | 2, 3, 24-31 | AvgCharge, Iept, Tiny, Small, Aliphatic, Non-Polar, Polar, Charged, Basic, Acidic | 10 |

The residue composition category includes (1) 20 features corresponding to the occurrence frequencies of each protein residue calculated for the whole query protein or its 25 residue-long C‐ or N-termini, and (2) 400 features corresponding to the normalized dipeptide residue composition of the whole protein or its termini. For the category of sequence/structure features, we include protein length and five other features calculated based on the predicted structural properties of the protein. The first three features are defined as the number of residues classified to be a part of *α*-helix, *β*-sheet, or loop secondary structures, respectively. The secondary structure is predicted using the SSpro/ACCpro package [30]. The other two features are defined as calculated percentages of exposed and buried residues in the protein, respectively. They are calculated based on the predicted relative solvent accessibility obtained using the same software package.

Physico-chemical properties are expected to play important roles in identifying effector proteins. For instance, the isoelectric point may be an important property since effector proteins have to exist and function in both the host and pathogenic environments. Thus, our final category includes ten physicochemical features defined as the percentage of residues of each of the seven physico-chemical types: acidic/basic, polar/non-polar, charged, tiny, and aliphatic [31], as well as the average charge and isoelectric point values. The physico-chemical types are calculated using EMBOSS [32].

### Analysis of feature importance, feature selection, and model training

The importance of the features may vary for each of the three groups of models. To rank the features by their importance and select a subset of important features for each group, we use the correlation-based feature selection (CFS) method [33]. The CFS method evaluates the importance of features based on the property that the subsets of important features are highly correlated with the class and are not correlated with each other. In contrast to a basic “greedy” method, where the importance of features is estimated through one-by-one feature removal, CFS allows to independently evaluate subsets of features. As a result, the features have been ranked, and the subsets of 27, 28, and 51 features are selected to train the N-terminal, C-terminal, and full sequence classifiers, respectively (Supplementary Table S3).

During SVM model training, a kernel type is selected for each model that maximizes the model’s f-measure. The models are trained and tested using the *libsvm* software package [34]. Our key goal is to minimize the number of proteins erroneously misclassified as effectors, *i.e*. false positives, while trying to maximize the number of predicted real effectors. We next explore whether this can be achieved through more stringent classification criteria defined next. Given an SVM model *M* and a training data of size *n*, for each training example *x_k_* let *f_k_* ∈ [+1, −1] be its decision value predicted by the SVM model, and *y_k_* ∈ [+1, −1] to be its true annotation of being an effector or non-effector. For each of the three SVM models, *M^(i)^*, the prediction probability for a training example *x_k_* is defined as 
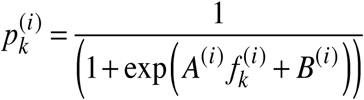
. The coefficients *A^(i)^* and *B^(i)^* are estimated during the SVM training process by minimizing the log-likelihood function. While the original prediction probability uses the probability threshold *θ*_0_ = 0.5 to classify the example as effector or non-effector, the optimal probability threshold is found using a simple grid search with the threshold step *Δθ* = 0.05.

Finally, to explore another state-of-the-art supervised classifier that is independent of a threshold, we train three Random Forest (RF) models using the same three subsets of selected features. The scikit-learn Python library was used for the training with the default parameters [35].

### Integrating three independent classifiers

Having trained three independent classifiers, we next would like to see if integrating the prediction results from all three classifiers can increase the prediction accuracy. To do so, we combine the independent SVM predictions for each protein into a single classifier using three simple integration schemes. The first scheme is a *minority voting*, in which a protein is classified as effector if at least one of three classifiers labels it as effector. The second scheme is *majority voting* in which the label corresponds to the predominant label of three classifiers. The last scheme is *unanimous voting*, in which a protein is labeled as effector if all three classifiers agree. For each voting scheme, all 27 combinations of SVM models over three SVM kernels are explored (Supplementary Table S4). The performance of each of the integration schemes is compared against the top performing independent classifier. We hypothesize that at least one of the integration schemes will perform better than any of the individual classifiers, if all independent classifiers demonstrate comparable performance.

### Method evaluation

#### Assessment of PREFFECTOR classifiers

For the SVM classifiers, the assessment of the methods is done after each of the three training stages: training the individual SVM models, optimization of the individual SVM models through feature selection, and integrating the three SVM models into a single classifier. Specifically, a standard validation on a previously unseen testing set and two cross validation protocols are implemented, including (1) 10-fold cross validation for all models, and (2) leave-one-out (LOO) cross validation for the top performing classifier (the latter protocol is time consuming hence its limited application). The choice of cross validation protocols, in addition to using a simple training set/testing set assessment scheme, is due to the limited number of effectors (and non-effectors, because of the balanced positive and negative subsets), which prevents having a large testing set. In addition, the assessment of the individual RF models is done using the same two cross validation protocols. For a more detailed explanation, see *Training sets* subsection of *Results* section.

In the first cross validation protocol, we generate the classification models using a 90%-10% split of the entire dataset, where 90% is used for training and 10% is used for testing. To alleviate any bias towards selecting a specific dataset for training and testing, both datasets are selected randomly, but with an additional constraint: we require that the positive training and testing datasets include similar numbers of effectors from all six secretion systems (Supplementary Tables S1, S2, S5). The dataset is shuffled between each of ten folds, where the positive training and testing datasets as well as negative training and testing datasets are randomly selected to minimize any overlap between the folds. The four evaluation measures (see below) are calculated for each of the ten folds, and their averages are reported as the final assessment. The LOO is an *N*-fold cross validation, where *N* is the size of the dataset. We note that for each fold of the 10-fold or LOO cross validations, the training and testing sets do not overlap.

For each set of SVM models, we evaluate the results using three different kernels: linear, polynomial, and radial basis function (RBF). Each classification task performed by both SVM and RF classifiers is evaluated using accuracy (*Acc*), recall (also called sensitivity, *Rec*), precision (*Pre*), and *f-measure*:

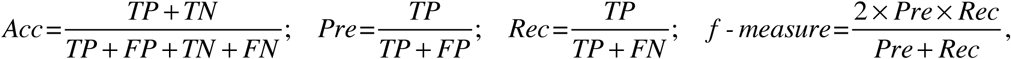

where *TP* is the number of true positives (effectors), *TN* is the number of true negatives (non-effectors), *FP* is the number of false positives, and *FN* is the number of false negatives. While each of the above four measures can be used to evaluate the overall performance of an effector prediction method, the recall measure can be used to evaluate the accuracy of predicting the effectors from a specific secretion system, which was also used in our comparative analysis of the state-of-the-art effector prediction methods.

#### Comparison with the state-of-the-art single secretion system prediction methods

We next would like to test if any of the existing state-of-the-art methods that are trained to predict effectors from a single secretion system can be applied to accurately predict effectors from the other secretion systems. To test this, we apply the existing methods to the same dataset of effectors as was used in LOO and 10-fold cross validation, calculating for each method the six recall values that correspond to the method’s predictions for each secretion system. We note that the other evaluation measures considered above depend on the negative set, which is not secretion system specific. Therefore, one cannot obtain the secretion system specific accuracy, precision, or *f*-measure. Nevertheless, the overall performance of the state-of-the-art methods was obtained for all four measures and compared with the performance of PREFFECTOR.

### GO function enrichment and depletion analysis

When applying PREFFECTOR on a whole-genome scale, the first question that one can ask is whether the predicted effectors share any common functions. We use the gene ontology (GO) annotation to functionally characterize each predicted effector and to analyze the functional prevalence among the effectors. Specifically, we employ the GOanna tool, developed as a part of AgBase resource [36], which maps GO terms based on sequence similarity to the previously annotated sequences. All obtained GO terms are then mapped to the second highest level in the GO hierarchy using CateGOrizer [37]. As a result, each effector is assigned a first-level term (biological process (P), molecular function (F), or cellular component (C)) and one or several of 60 unique second-level terms, when possible (Supplementary Table S6). Selection of the GO annotation level is a trade-off between the annotation sparseness across the genes and the level of details that a GO term provides: the first level provides too general terms, while selecting the third level has lead to very specific annotations of individual genes for which no statistical analysis is possible. It is important to note that no GO functional annotation was used for the input features in the classification task, since we did not want to limit the coverage of PREFFECTOR by excluding novel genes with no functional annotation available.

The GO function depletion/enrichment analysis is performed for the set of effectors from each analyzed bacterial genome. We use the two-sided mid-*P*-value doubling approach [38]. Specifically, given a function F defined by a GO category, our mid-*P*-value is calculated as:

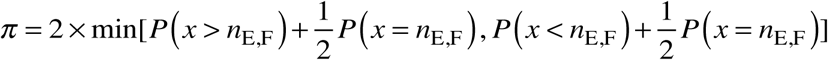

where:

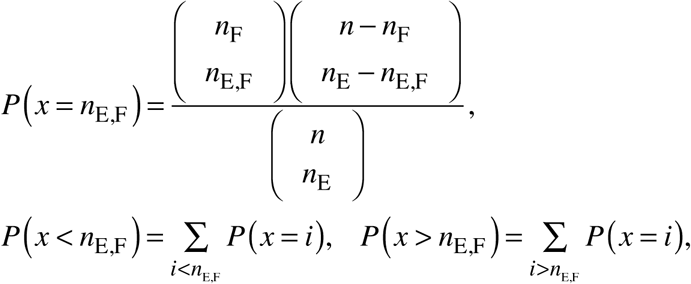

and *n*_E,F_ is the number of effectors in the genome that are annotated with function F, *n*_F_ is the total number of genes annotated with F, *n*_E_ is the total number of effectors, and *n* is the total number of genes in the genome. In our analysis, the significance level is considered to be *α* = 5%. The doubling approach while being comparable with the minimum-likelihood approach is computationally less expensive than the latter. In addition, the use of mid-P-value allows minimizing the loss of power due to the discreteness of the underlying hypergeometric distribution.

## Results

### Training sets

The positive set of experimentally verified effectors was collected and manually curated by combining the following two search strategies. In the first strategy, we retrieved bacterial effectors by mining the UniProt database [39] using the keyword-based search followed by manual curation of sequences. During the manual curation, only those sequences were considered that had been annotated and had a publication reference with experimental evidence of the translocation into the host cells through a secretion system. In the second strategy, we directly searched the PubMed database using keywords associated with protein effectors, followed by manual curation of the retrieved information to ensure that each effector is supported by experimental evidence from at least one publication. We then merged both sets obtained from the two strategies, removing all identical sequences or sequences from different strains of the same bacterial species. The obtained curated set of 323 effectors (Supplementary Materials) is diverse species-wise (78 unique bacterial species and strains) and protein-wise (protein lengths ranging from 56 to 5,559 residues). To the best of our knowledge, this is the largest curated set of effectors that covers all six secretion system types. Due to the prevalence of data on well-studied systems, such as T3SS, we further reduced the set of effectors, making the final set to be more evenly distributed across all six types of secretion systems. At the same time, the homology-based sequence redundancy was removed such that pairs of sequences in the resulting dataset did not share high sequence identity, according to a standard Needleman-Wunsch global alignment (<u>*nwalign*</u> tool of MATLAB was used). In the final dataset of 168 effectors, 160 proteins do not share more than 30% sequence identity with one another; 6 other proteins share less than 40% sequence identity with at least one more protein, and 2 other proteins share less than 50% sequence identity. The resulting positive set included 168 effectors (Supplementary Tables S1, S5).

To collect the negative data set of non-effectors, we used another UniProt-based strategy. For each organism that contributed an effector protein, we retrieved its proteome from UniProt and removed all protein sequences that occurred in the positive set or any other protein that was annotated by UniProt as an effector. Then we selected proteins that cover a wide range of housekeeping functions and are spread across different parts of the cell. The manually curated negative set of 168 non-effectors includes proteins from 70 bacterial species, with the proteins lengths ranging between 56 and 2,703 residues. Similar to the positive data set, proteins in the negative data set do not share close homology: none of 168 non-effectors, share more than 30% sequence identity with one another, based on the Needleman-Wunsch global alignment.

The final dataset consists of a total of 336 proteins: 168 effector proteins in the positive set and 168 non-effectors in the negative set, obtained from the same species as effectors (Supplementary Table S2).

### Feature analysis reveals important features shared between all classifiers

The most important features for each of the three groups of classifiers were selected and analyzed using the correlation-based feature selection (CFS) approach [33]. We note that the CFS algorithm does not select a predefined number of features. As a result, all three groups of classifiers had different numbers of selected important features: 20 for N-terminal model, 23 for C-terminal model, and 49 for the full model. While the overall sets of selected features differed significantly between the three classifiers, several features were found commonly important (Supplementary Table S3).

Some of the found commonly important features have been supported by the previous findings. For instance, all three models reported as important the occurrence frequencies for the two individual residues, serine and asparagine, as well as more general, small and charged, residue types [31]. Indeed, the enrichment of serine residues in the N-termini have been observed as common features in the effectors of organisms hosting T3SS [22]. Conserved serine positions have been reported as a requirement for the proteins’ enzymatic activity of a family of bacterial effectors targeting plants [40]. Furthermore, it has been reported that many effectors have membrane localization domains (MLDS) that are enriched in charged residues [41]. Lastly the importance of a feature characterizing the presence of a coil, or loop, secondary structure type in the effector sequence can be supported by the previously reported observations of a high abundance of intrinsically disordered regions in effectors [42]. Other common features selected by the CFS method, such as occurrence frequencies of asparagine and small residues have not been previously reported.

### Method assessment and comparison with existing tools

#### Performance assessment

The assessment of the individual classifiers based on the C-terminal, N-terminal, and full sequence RF and SVM models has revealed an unexpected but consistent difference between the models’ accuracies (Table 3, Supplementary Tables S7, S8, S9). Specifically, there is a significant gap in performance when comparing the full models with C‐ and N-terminal models: the LOO accuracies of the top performing full, N-, and C-terminal models were 0.89, 0.71, and 0.74, respectively. The top performing full model was RF classifier; the full SVM model with the RBF kernel also achieved similar accuracy. Interestingly, even the worst performing full model (polynomial kernel) was more accurate than the best N‐ and C-terminal models (Table 3). The most accurate full model was also shown to be consistent in its performance across all six types of secretion systems (Figure 1C).

**Table 3.** Leave-one-out (LOO) assessment for three SVM and Random Forest (RF) models. The performance of RF classifier and each of three SVM classifiers using one of the three different kernels, Radial Base Function (Radial), Linear, and Polynomial, is measured by Accuracy (Acc), Precision (Pre), Recall (Rec), and f-measure. The default probability threshold of *θ*_0_ = 0.5 is used for each SVM model.

| ***Model*** | ***Kernel*** | ***Acc*** | ***Pre*** | ***Rec*** | ***f-measure*** |
| --- | --- | --- | --- | --- | --- |
| N | Radial | 0.80 | 0.80 | 0.80 | 0.80 |
|  | Linear | 0.76 | 0.77 | 0.76 | 0.76 |
|  | Polynomial | 0.68 | 0.73 | 0.68 | 0.67 |
|  | RF | 0.71 | 0.71 | 0.71 | 0.71 |
| C | Radial | 0.73 | 0.73 | 0.73 | 0.73 |
|  | Linear | 0.70 | 0.70 | 0.70 | 0.70 |
|  | Polynomial | 0.67 | 0.73 | 0.67 | 0.64 |
|  | RF | 0.68 | 0.68 | 0.68 | 0.68 |
| F | Radial | 0.87 | 0.87 | 0.87 | 0.87 |
|  | Linear | 0.83 | 0.83 | 0.83 | 0.83 |
|  | Polynomial | 0.81 | 0.84 | 0.81 | 0.81 |
|  | RF | 0.89 | 0.89 | 0.89 | 0.89 |

Next, by performing the grid search based optimization of full models with the polynomial and RBF kernels, we explored if the false positive rate, FPR, could be significantly lowered while preserving the overall accuracy (Table 4, Supplementary Table S10). In both cases, a slightly higher probability threshold of *θ*_0_ = 0.4 resulted in a slightly higher accuracy for both models with polynomial and RBF kernels, while the false positive rate, FPR, also increased. On the other hand, even the maximized accuracy for either kernels of the SVM model was worse than the accuracy (and f-measure) for the RF classifier. As a result, the full model using Random Forest classifier was implemented as a default classifier in our PREFFECTOR web-server. Furthermore, a threshold of *θ*_0_ = 0.9 and 0.8 for the RBF and polynomial kernels drastically reduced the FPR to 0.03 while reducing the accuracy to 0.74. Thus, to further minimize the number of false-positives due to a disproportionally high number of non-effectors when applying our method on the whole-genome scale, we have implemented in the web-server an SVM model that uses the RBF kernel and a stringent threshold of *θ*_0_ = 0.9 as an alternative to the default model when using it on a large set of proteins from the same genome.

**Table 4.** Grid search of the probability threshold *θ*_0_ **for a full SVM model with RBF kernel and 10-fold cross validation assessment.** While the original prediction probability uses the probability threshold *θ*_0_ = 0.5, the optimal, with respect to the accuracy probability thresholds of *θ*_0_ = 0.4 and 0.3 increase the false positive ratio and are not considered. In turn, the most stringent threshold of *θ*_0_ = 0.9 drastically decreases the false positive rate, while decreasing accuracy by 0.11. Abbreviations: TP – true positives, TN – true negatives, FP – false positives, FN – false negatives, FPR – false positive rate, TPR – true positive rate.

| $\theta$ | TP | TN | FP | FN | FPR | TPR | Accuracy |
| --- | --- | --- | --- | --- | --- | --- | --- |
| 0.1 | 15 | 8 | 8 | 1 | 0.48 | 0.93 | 0.73 |
| 0.2 | 15 | 12 | 5 | 1 | 0.29 | 0.91 | 0.81 |
| 0.3 | 15 | 14 | 2 | 1 | 0.17 | 0.89 | 0.86 |
| 0.4 | 14 | 14 | 2 | 2 | 0.14 | 0.86 | 0.86 |
| 0.5 | 13 | 15 | 2 | 3 | 0.12 | 0.82 | 0.85 |
| 0.6 | 13 | 15 | 1 | 3 | 0.11 | 0.78 | 0.84 |
| 0.7 | 12 | 15 | 1 | 4 | 0.07 | 0.72 | 0.83 |
| 0.8 | 10 | 16 | 0 | 6 | 0.05 | 0.64 | 0.79 |
| 0.9 | 8 | 16 | 0 | 8 | 0.04 | 0.51 | 0.74 |

The analysis of each of the three integration schemes revealed that neither of the schemes could improve the accuracy of the top-performing individual classifiers (full sequence model with polynomial and RBF kernels). Overall, the majority voting performed significantly better than the other two voting schemes (Supplementary Table S4). Due to its inferior performance at the evaluation stage, no integration model was included into our web-server.

#### Comparison with state-of-the-art methods

Finally, we tested whether any of the previously published methods, which were designed to determine effectors in a specific secretion system, could be applied to other secretion systems as well. We hypothesized that the features used by those classifiers could be sufficient to determine effectors in the other secretion systems. We ran seven top-performing effector classifiers from Table 1 for which we could access a stand-alone tool or a web-server to our dataset used in LOO cross-validation. The results showed, as expected, that none of the currently existing effector predictors could be used as a broad effector classifier (Fig.1D, Supplementary Tables S11, S12), with the overall accuracies ranging from 0.45 to 0.66 and recall values ranging from 0.04 to 0.48. The evaluation of these methods across different secretion systems has confirmed the utility of each algorithm in predicting effectors for one particular secretion system. Interestingly, for many predictors their performance on our dataset was worse than in the original publications. This could be attributed to the fact that our dataset was designed to exclude the majority of homologous proteins, which could potentially bias the prediction accuracy. Another reason for the better performance of PREFFECTOR could be the inclusion of new effectors into its training set, with some of the effectors carrying a novel secretion signal.

### Application to whole-genome search for effectors in bacterial pathogens

Our method presents an efficient way for zeroing-in on putative effector candidates on the genome scale. To illustrate this, we have applied our method to four genomes of Gram-negative bacteria, *Acinetobacter baumannii*, *Chlamydia trachomatis*, *Helicobacter pylori*, and *Legionella pneumophila* (Fig. 2, Supplementary Table S13, and Supplementary Figs. S1-S3). The four bacteria are infectious agents of severe infectious diseases with limited knowledge on virulence factors or secreted effectors.

**Figure 2.**
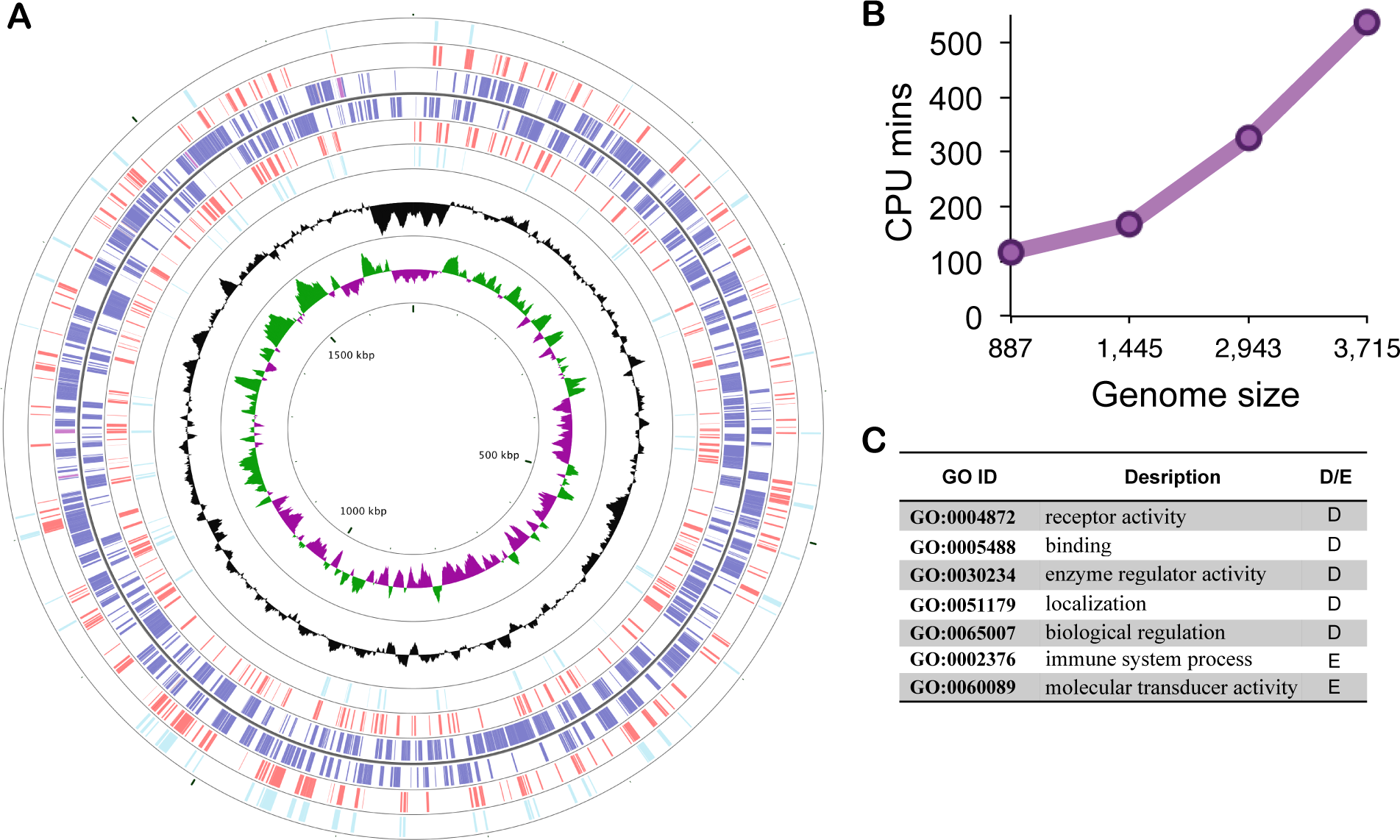
Whole-genome application of PREFFECTOR. A. Predicted effectors of Chlamydia trachomatis using the optimal probability threshold of *θ*_0_ = 0.6 (red) and a stringent probability threshold of *θ*_0_ = 0.99 (cyan) are mapped according to their corresponding positions on the circular bacterial genome. Many predicted effectors for both thresholds form distinct compact clusters. All known genes (blue) are mapped on the corresponding DNA strand of the genome. Shown in black is GC content. Shown in green and purple are GC skew+ and GC skew-, respectively. Regions of tightly clustered effectors can be clearly identified. The image was generated using CGView Comparison Tool. B. The analysis of the PREFFECTOR’s performance on the four whole genomes (*Acinetobacter baumanni, Chlamydia trachoma, Helicobacter pylori, and Legionella pneumophila*). C. Examples of enriched (E) and depleted (D) GO functions occurring in the three genomes; both enriched are depleted GO functional terms are unique to each genome.

*Acinetobacter baumannii* is a member of ESKAPE (*Enterococcus faecium, Staphylococcus aureus, Klebsiella pneumoniae, Acinetobacter baumannii, Pseudomonas aeruginosa* and *Enterobacter* species) pathogen group of bacteria with a high rate of antimicrobial drug resistance and causing nosocomial infections [43, 44]. Mortality in the patients infected by *A. baumannii* is reported to be between 30% and 75% [45]. The genome has been shown to encode a T4SS, and more recently type VI secretion system (T6SS), however a comprehensive list of effectors secreted by each apparatus is yet to be found [46, 47]. The genome size is roughly 4 Mb and varies in size depending on the strain [48]. In this work, the genome of 1656_2 strain was used, which includes 3,715 genes. The application of RF-based PREFFECTOR to this genome resulted in 753 predicted effectors (Supplementary Fig. S1). Interestingly, all of the only five previously experimentally identified effectors were correctly labeled by our method.

*Chlamydia trachomatis* is a cause for the world’s most prevalent bacterial sexually transmitted infection, with a global estimate of 130 millions new cases in 2012 [49]. While no high-throughput study to identify effectors in *C. trachomatis* has been done to date, more than a dozen effectors have been identified and experimentally validated [50, 51]. The genome of this Gram-negative bacterial species is around 1 Mb long and hosts T3SS. The specific strain used in our study includes 926 genes. The application of RF-based PREFFECTOR found 198 genes annotated as effectors, including 12 (57%) of the 21 known effectors (Fig. 2A).

*Helicobacter pylori*, one of the world’s most common pathogenic bacteria has been associated with chronic gastritis, gastric ulcers, stomach cancer, and other diseases [52-54]. The bacterial genome size is 1.6-1.7 Mb, depending on the strain [55]. The most well studied pathogenicity island, *cag*, is known to carry type IV and V secretion system machinery [56-58]. However, only a few *H. pylori* effectors have been identified to date [28]. Our RF-based prediction has identified 457 effectors out of 1,485 genes identified in the genome of J99 strain of the bacteria, including all three previously known effectors (Supplementary Fig. S2).

*Legionella pneumophila* is the agent of Legionnaires’ disease, a form of atypical pneumonia with the estimated fatality rate of almost 30% [59], occurring primarily in the undeveloped countries, but with the most recent 2015 outbreaks in the New York City and Northern California. Out of 180 known *Legionella* strains, only a few of them have been fully sequenced [60, 61]. The size of the sequenced genomes ranges between 3.3 and 3.6 Mb and contains ~3,000 genes. Effectors of *L. pneumophila* are secreted through secretion systems of types II and IV, and possibly I and V [62]. Our RF-based PREFFECTOR classifier predicted 989 effectors out of known 3,025 genes of Paris strain of *L. pneumophila* [63], including 105 (78%) of 134 known effectors (Supplementary Fig. S3).

Last, when applying the stringent threshold of *θ*_0_ = 0.9 to *Acinetobacter baumannii*, *Chlamydia trachomatis*, *Helicobacter pylori*, and *Legionella pneumophila*, the obtained numbers of effectors were significantly smaller: 260, 72, 119, and 238 effectors, correspondingly.

The GO functional enrichment analysis did not find common functions that are significantly enriched or depleted across all three species; the fourth species, *Chlamydia*, did not result in any significantly enriched or depleted GO terms.. The predicted effectors had primarily significantly depleted functions, including biological regulation, binding, localization, and enzyme regulator activity (Fig 2C, Table 5, Supplementary Table S6). The only two significantly enriched functions, immune system process and molecular transducer activity, were found in *Acinetobacter.* We recall that no GO functional annotation was used for the input features of PREFFECTOR classifiers.

**Table 5.** Significantly enriched (E) and depleted (D) GO functions for the three bacterial genomes. There were no significantly enriched and depleted GO functions found in *Chlamydia*. Effectors were predicted using Random Forrest classifier.

| *Acinetobacter* |  |  | *Helicobacter* |  |  | *Legionella* |  |  |
| --- | --- | --- | --- | --- | --- | --- | --- | --- |
| *GOTerm* | *p-value* | *E/D* | *GOTerm* | *p-value* | *E/D* | *GOTerm* | *p-value* | *E/D* |
| GO:0065007 | 1.7E-03 | D | GO:0051179 | 1.3E-02 | D | GO:0030234 | 4.4E-02 | D |
| GO:0005488 | 5.4E-03 | D | GO:0065007 | 2.0E-02 | D |  |  |  |
| GO:0005623 | 6.0E-03 | D | GO:0004872 | 5.3E-02 | D |  |  |  |
| GO:0003824 | 2.2E-02 | D |  |  |  |  |  |  |
| GO:0002376 | 4.1E-02 | E |  |  |  |  |  |  |
| GO:0060089 | 5.5E-02 | E |  |  |  |  |  |  |

The performance time of our algorithm for all four genomes was consistent, taking ~2 min to classify a gene, or between two and nine hours per a genome when running it on a 15-core computing server with Intel Xeon 2.3 GHz CPU processors (Fig. 2B). We note that the time it takes to process a single protein sequence depends on the length of the protein sequence and may vary significantly. The sets of predicted effectors, ranked by the prediction confidence, for each of the four genomes are available for download at the PREFFECTOR website.

### PREFFECTOR web-server

We have implemented the top-performing classifiers as PrEffector web-server that allows batch submission of up to 500 protein sequences per each job. The web-server features very simple input and output interfaces. As the only input parameter, a user is asked to select a default mode (corresponding to Random Forest classifier) or the one that drastically reduces the number of false positives, leaving in only the near-certain predictions (SVM with RBF kernel and threshold of *θ*_0_ = 0.9). The output includes the FASTA header of each input sequence, its classification as effector or non-effector and prediction’s probability according to the implemented classifier. When the job is submitted, the user is provided with the information on the job’s position in the queue and the unique job reference number. Once the job is completed, the user is notified via email.

## Discussion

### PREFFECTOR is complimentary to methods predicting effectors from a single secretion system

In this work, we have presented a ready-to-use sequence-based computational tool to predict bacterial effectors in the genomes of Gram-negative bacteria, irrespective of secretion system type(s) they carry. Recently, a number of accurate secretion system-specific methods have been presented, targeting mainly T3SS and T4SS. The top performing methods report the accuracy in the 90 percent range, presenting a clear selection choice when one knows that the pathogen carries one of the two secretion systems, and only that system. Unfortunately, the lack of complete knowledge about effectors and their secretion systems for a vast number of bacterial genomes make applications of such specialized methods limited. In addition, our assessment of the top performing methods on an independent dataset that is balanced across the six secretion systems has shown that, as expected, none of the tested specialized methods are generalizable for an arbitrary secretion system. Thus, we consider our approach as truly complementary to the existing secretion system-specific methods, and perhaps the first method to employ when a new bacterial genome is sequenced but no prior knowledge about its secretion system(s) is available. This possibility of the *ab-initio* discovery of novel effectors—when the knowledge of secretion signals (or secretion system in general) is lacking, or when there is more than one secretion system in the genome—is the main advantage of our approach.

A comprehensive assessment of our method showed consistent performance of the most accurate, full sequence SVM model, with high and nearly identical accuracy, precision, and recall values. It has also revealed that two other models (N-terminal and C-terminal SVM models, respectively) are significantly inferior to the top-performing model. One possible reason is the fact that different effectors house their secretion signals in different parts of their sequences; for many effectors the location (or even existence) of the secretion signal is unknown. Thus, focusing on just a part of the protein sequence, *e.g.* its C-terminus, may lead to loosing critical information for some proteins that have the secretion signal located in the other part of the sequence, *e.g.* N-terminus. In addition, the poor performance of the latter two models is likely the key contributing factor to the lack of improvement in the prediction accuracy when using any of the three integration schemes.

A possible venue for improving the accuracy of PREFFECTOR is to train it on an unbalanced dataset of effectors and non-effectors that reflects the real percentage of effectors in the whole bacterial genome. Our preliminary experiments with different effectors-to-non-effectors ratios demonstrated significantly reduced accuracy, and therefore we decided not to pursuit this direction further, instead focusing on the balanced training set. Implementing a more stringent classification threshold was suggested as an alternative solution that reduced the number of false positives while increasing the number of false negatives.

### Resolving issues with the whole-genome application of PREFFECTOR

The application of PREFFECTOR to four bacterial genomes has demonstrated the feasibility of our method to predict effectors on a whole-genome scale, allowing for large-scale comparative studies of bacterial effectors. Yet, the application has revealed that our method is yet to achieve the perfect performance: the number of predicted effectors using the optimized classifier threshold (*θ*_0_ = 0.6) appeared to be high, and also missed some experimentally validated effectors. Recall that we could only formally assess the accuracy of our method based on the three rigorous assessment protocols and by comparing its performance with other methods on the same dataset, in which our approach outperformed every single method for every single secretion system. However, we also applied the top performing methods specialized in predicting effectors of T3SS and T4SS to *C. trachomatis*, *H. pylori*, and *L. pneumophila,* which are known to host at least one of the two secretion systems. The results showed that with *θ*_0_ = 0.6, our method predicted a comparable number of the effectors in *C. trachomatis*, while the numbers of predicted effectors in *H. pylori* and *L*. *pneumophila* by our method were higher than by other classifiers (see Table S14). While some of the predicted effectors would unavoidably be the false positives according to our assessment, there are several factors that could support the high numbers of effectors obtained by our method. First, both *H. pylori* and *L. pneumophila* are known to host multiple secretion systems. So, it is likely that PREFFECTOR identifies effectors from other secretion systems, which cannot be predicted by T4SS-specialized methods. Second, approximately 300 effectors have been already experimentally identified in *L*. *pneumophila* [63] including 134 obtained from a single screening experiment [7]. The large number of effectors may be attributed to the functional redundancy where multiple effectors perform similar functions and target different steps of the same molecular pathway in the host. The option of a stringent threshold (*θ*_0_ = 0.9) implemented in the PREFFECTOR web-server allows one to zero-in on a small number of the most likely effector candidates that can be confirmed experimentally.

With the growing number of sequenced genomes of pathogenic bacteria, many of which are hardly studied, the importance of a free, fast, and accurate tool for in-silico prediction of the effector candidates cannot be overstated. The developed PREFFECTOR classifier allows moving one step further in this direction. We expect our method to impact research on emerging and neglected diseases by guiding effector targeting experiments and antibiotics design, and making such studies faster.

## Acknowledgements

We thank Anitha Subramani for useful discussions and help during the initial development of PREFFECTOR. This work has been supported by National Science Foundation (DBI-0845196 to DK), and by Agriculture and Food Research Initiative Competitive Grant no. 2015-67013-23511 from the USDA National Institute of Food and Agriculture.

## Author Contributions Statement

DK conceived the idea. AD, SW, and DK collected and curated the dataset of effectors. AD implemented PREFFECTOR method and web-server. SW performed GO functional analysis. AD and DK analyzed the data. All authors contributed to the writing of the manuscript.

## Supporting Information Legends

**Figure S1 – Predicted effectors of *Acinetobacter baumannii*.** Effectors were predicted using the default optimal model (Random Forest, red) and SVM with extremely stringent threshold (*θ*=0.9, cyan) are mapped according to their corresponding positions on the circular bacterial genome. All known genes (blue) are mapped on the corresponding DNA strand of the genome. Shown in black is GC content. Shown in green and purple are GC skew+ and GC skew-, respectively. Regions of tightly clustered effectors can be clearly identified. The image was generated using CGView Comparison Tool.

**Figure S2 – Predicted effectors of *Helicobacter pylori*.** Effectors were predicted using the default optimal model (Random Forest, red) and SVM with extremely stringent threshold (*θ*=0.9, cyan) are mapped according to their corresponding positions on the circular bacterial genome. All known genes (blue) are mapped on the corresponding DNA strand of the genome. Shown in black is GC content. Shown in green and purple are GC skew+ and GC skew-, respectively. Regions of tightly clustered effectors can be clearly identified. The image was generated using CGView Comparison Tool.

**Figure S3 – Predicted effectors of *Legionella pneumophila*.** Effectors were predicted using the default optimal model (Random Forest, red) and SVM with extremely stringent threshold (*θ*=0.9, cyan) are mapped according to their corresponding positions on the circular bacterial genome. All known genes (blue) are mapped on the corresponding DNA strand of the genome. Shown in black is GC content. Shown in green and purple are GC skew+ and GC skew-, respectively. Regions of tightly clustered effectors can be clearly identified. The image was generated using CGView Comparison Tool.

**Table S1. Distribution of effectors in the positive set across six secretion system types.** To exclude bias, the effectors are evenly distributed across the secretion systems and are selected from different species.

**Table S2. Distribution of positive (effectors) and negative (non-effectors) sets across different genera.** Reported for each genus are the numbers of effectors, non-effectors, and secretion systems that those effectors belong to.

**Table S3. Results of feature selection for each SVM model.** Features that were found commonly important across the three models are underscored and in bold.

**Table S4. Assessment of three integration schemes.** Shown are the accuracy values for each voting scheme. All 27 combinations of SVM models over three SVM kernels are explored.

**Table S5. A detailed list of effectors in the positive set.** Each effector is included together with the species it belongs to and the secretion system it is identified with.

**Table S6. GO annotation terms used in functional enrichment analysis.** Shown is the list of second-level terms used in the analysis.

**Table S7. Leave-one-out (LOO), 10–fold cross validation (CV), and test set assessments for N-terminal SVM and Random Forest (RF) models.** For each assessment type, the performance using one of the three different kernels is measured by Accuracy (Acc), Precision (Pre), Recall (Rec), and f-measure. The default probability threshold of *θ*_0_ = 0.5 is used.

**Table S8. Leave-one-out (LOO), 10–fold cross validation (CV), and test set assessments for C-terminal SVM and Random Forest (RF) models.** For each assessment type, the performance using one of the three different kernels is measured by Accuracy (Acc), Precision (Pre), Recall (Rec), and f-measure. The default probability threshold of *θ*_0_ = 0.5 is used.

**Table S9. Leave-one-out (LOO), 10–fold cross validation (CV), and test set assessments for the full sequence SVM and Random Forest (RF) models.** For each assessment type, the performance using one of the three different kernels is measured by Accuracy (Acc), Precision (Pre), Recall (Rec), and f-measure. The default probability threshold of *θ*_0_ = 0.5 is used.

**Table S10. Grid search of the probability threshold** *θ*_0_ **for a full SVM model with RBF kernel and 10-fold cross validation assessment.** While the original prediction probability uses the probability threshold *θ*_0_ = 0.5, the optimal, with respect to the accuracy probability threshold of *θ*_0_ = 0.4 increases the false positive ratio and is not considered. In turn, the most stringent threshold of *θ*_0_ = 0.9 drastically decreases the false positive rate, while significantly decreasing accuracy by 0.16. Abbreviations: TP – true positives, TN – true negatives, FP – false positives, FN – false negatives, FPR – false positive rate, TPR – true positive rate.

**Table S11. Comparative performance analysis of sate-of-the-art effector prediction methods on our dataset.** The application of the methods that are designed to detect effectors of a single secretion system on the dataset of 168 effectors and 168 non-effectors has showed that none of them can be used universally across all six secretion systems.

**Table S12. Recall values across six secretion systems for each state-of-the-art method applied to our dataset.** One asterisk corresponds to the methods specialized in predicting T3SS effectors, while two asterisks correspond to the methods specialized in predicting T4SS effectors.

**Table S13. Whole-genome application of PREFFECTOR to four Gram-negative bacterial species.** Shown are application results for the default optimal (Random Forest) and SVM with extremely stringent threshold (*θ*=0.9) predictions. The smaller number of known effectors for the classifier that uses the extremely stringent threshold is due to a higher number of false negatives.

**Table S14. Whole-genome application of specialized state-of-the-art prediction methods to the same four Gram-negative bacterial species.** Shown are the numbers of genes predicted as effectors. Only stand-alone software packages were used in this application.

**Supplementary Data.** Complete List of Effectors.

